# Pinging the Brain with Transcranial Magnetic Stimulation Reveals Cortical Reactivity in Time and Space

**DOI:** 10.1101/2019.12.18.880989

**Authors:** Sangtae Ahn, Flavio Fröhlich

## Abstract

Single-pulse transcranial magnetic stimulation (TMS) elicits an evoked electroencephalography (EEG) potential (TMS-evoked potential, TEP), which is interpreted as direct evidence of cortical reactivity to TMS. Thus, combining TMS with EEG may enable the mechanistic investigation of how TMS treatment paradigms engage network targets in the brain. However, there remains a central controversy about whether the TEP is a genuine marker of cortical reactivity to TMS or the TEP is contaminated by responses to peripheral somatosensory and auditory inputs. Resolving this controversy is of great significance for the field and will validate TMS as a tool to probe networks of interest in cognitive and clinical neuroscience. Here, we delineated the TEP’s cortical origins by localizing successive TEP components in time and space and modulating them subsequently with transcranial direct current stimulation (tDCS). We collected both motor evoked potentials (MEPs) and TEPs elicited by suprathreshold single-pulse TMS to the left primary motor cortex (M1). We found that the earliest TEP component (P25) was localized on the TMS target location (left M1) and the following TEP components (N45 and P60) largely were localized on the primary somatosensory cortex, which may reflect afferent input by hand-muscle twitches. The later TEP components (N100, P180, and N280) largely were localized to the auditory cortex. To casually test that these components reflect cortical and corticospinal excitability, we applied tDCS to the left M1. As hypothesized, we found that tDCS modulated cortical and corticospinal excitability selectively by modulating the pre-stimulus mu-rhythm oscillatory power. Together, our findings provide causal evidence that the early TEP components reflect cortical reactivity to TMS.

## Introduction

Combined transcranial magnetic stimulation (TMS) and electroencephalography (EEG) provide an opportunity to quantify brain network dynamics by pinging them with TMS[1]. The TMS-evoked potential (TEP), which is considered a reflection of cortical reactivity to TMS, has been shown to have diagnostic value in a variety of neurological and psychiatric disorders[2]. However, there is ongoing controversy about the origin of the TEP. A recent study claimed that the stimulation of peripheral nerves and the TMS coil’s loud clicking sound may confound the TEP amplitude[3]. Specifically, sham TMS elicited EEG potentials that were correlated highly with those by real TMS, despite the use of sophisticated procedures to attenuate the somatosensory and auditory confounds. In rebuttal of this publication, it was suggested that insufficient TMS intensity and incomplete auditory masking may explain the sensory-dominant evoked potentials in the experiment[4]. Nonetheless, residual auditory input is unavoidable in TMS studies[5] because of air and bone conduction from the TMS clicking sound[6,7]. Thus, it continues to be debated whether the TEP represents genuine cortical reactivity that single-pulse TMS elicits or whether it reflects cortical reactivity contaminated with peripherally- and auditory-evoked potentials. Here we sought to resolve this controversy and delineate TEPs by localizing the electrophysiological response with high-density EEG, structural magnetic resonance (MR) images, and digitized EEG electrode locations. If a TEP is localized in areas in the auditory and somatosensory cortex, then it can be determined that auditory input and peripheral nerve stimulation, respectively, drive this component. We chose the primary motor cortex (M1) as a stimulation target because the corticospinal response (motor-evoked potential, MEP) also should reflect cortical reactivity. To causally test the validity of our approach, we applied transcranial direct current stimulation (tDCS) to modulate cortical and corticospinal excitability[8,9]. We observed that single-pulse TMS to the M1 elicited six TEP components. The earliest TEP at 25ms (P25) from the TMS onset was localized to the hand area of the left M1, the TMS target location. The following two TEP components were localized to the primary somatosensory cortex (N45 and P60), which may reflect afferent input by hand-muscle twitches in response to suprathreshold TMS. The later TEP components (N100, P180, and N280) largely were localized to the auditory cortex. Further, tDCS modulated the cortical reactivity (TEP) and corticospinal response (MEP) reliably by modulating the pre-stimulus mu-rhythm oscillatory power. Together, our findings demonstrated that the earliest TEP component reflects genuine cortical reactivity, while the following TEP components may reflect different sensory processing.

## Results

### Cortical reactivity to single-pulse TMS

We investigated cortical reactivity to TMS in 18 healthy, right-handed, male participants (Fig. 1a). We obtained structural MR images (T1-weighted) for each participant using a 3T-MR scanner for precise targeting of the TMS with a neuronavigation system (Fig. 1b, left, Supplementary Fig. 1). After determining each participant’s resting motor threshold (RMT), we administered single-pulse TMS to the left M1 while recording TEPs and MEPs in three sessions. The tDCS condition (anodal, cathodal, and sham tDCS) was randomized for each session in a double-blind, cross-over study design. For tDCS, we used the conventional two-electrode montage (Fig. 1c, left, referred to as M1-SO, one for the M1, and another for the supraorbital cortex). We applied 100 single pulses of TMS before and after tDCS during each session. At the end of each session, we collected EEG electrode locations using a stereo-camera tracking digitizer to improve accuracy of source localization.

**Fig. 1.**
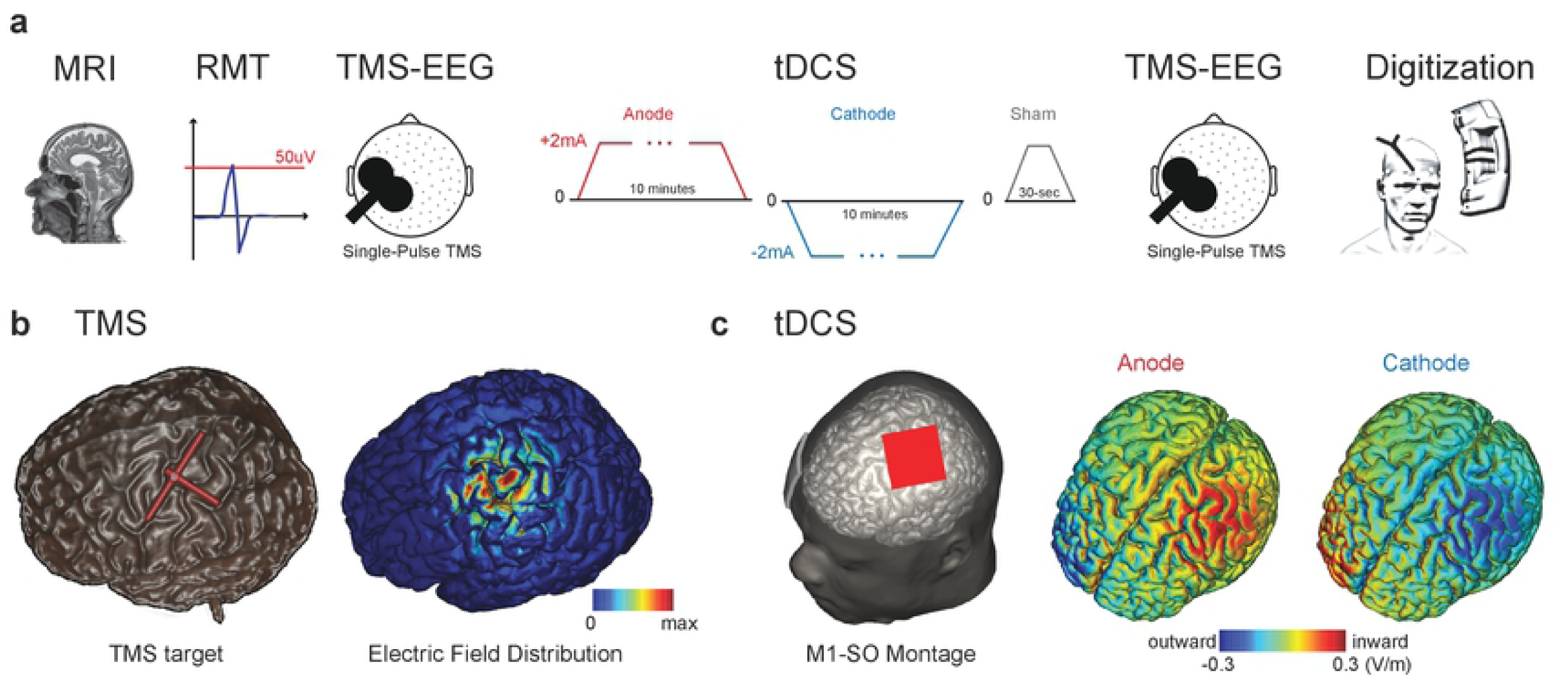
Experimental setup, stimulation setting, and electric field modeling of tDCS and TMS. (a) Structural MR images (T1-weighted) were obtained. The resting motor threshold was determined (MEP >50uV on 50% of the trials). Single-pulse TMS (100 pulses) was delivered to the hand area of the left M1 while recording TEPs and MEPs. Next, tDCS (anodal, cathodal, or sham tDCS) was applied for 10 minutes at 2mA current intensity with 30-sec ramp up and down periods. For sham tDCS, we applied 30 seconds of anodal tDCS as an active placebo. Each participant received all three tDCS conditions on a different day with at least a three-day interval between sessions. The order of the tDCS conditions was randomized and distributed equally across participants. Single-pulse TMS (100 pulses) was delivered after tDCS. Finally, EEG electrode locations were recorded by a stereo-camera tracking digitizer. (b) A representative example of the TMS target superimposed on the cortex (left). Red crosshair indicates the TMS coil’s position and orientation. Electric field distribution of TMS (right). (c) Carbon-silicone electrodes (5×5cm) were applied to the left motor hotspot (red square electrode) and supraorbital (SO) cortex (gray square electrode) referred to as the M1-SO montage (left). Inward and outward electric field distribution of anodal and cathodal tDCS (right).

Single-pulse TMS elicits multiple TEP components in the TMS-EEG recordings[10,11]. To determine whether single-pulse TMS to the hand area of the left M1 elicits TEPs, we computed grand-averaged TEPs (5255 epochs after bad epoch rejection) for each EEG channel from the TMS-EEG recordings before tDCS application. A butterfly plot of TEPs (Fig. 2a, gray lines) was obtained as a function of time (−100 to 500ms with respect to TMS onset) for each EEG channel (128 channels), and an averaged TEP over the left sensorimotor area (C3 channel and the 6 channels surrounding C3; see inset) was computed in sensor space (Fig. 2a, thick black line). We found that the averaged TEP on the left sensorimotor area exhibited three positive and three negative peaks relative to the baseline period (−100 to 0ms). We refer to these peaks by their canonical names: P25, N45, P60, N100, P180, and N280.

**Fig. 2.**
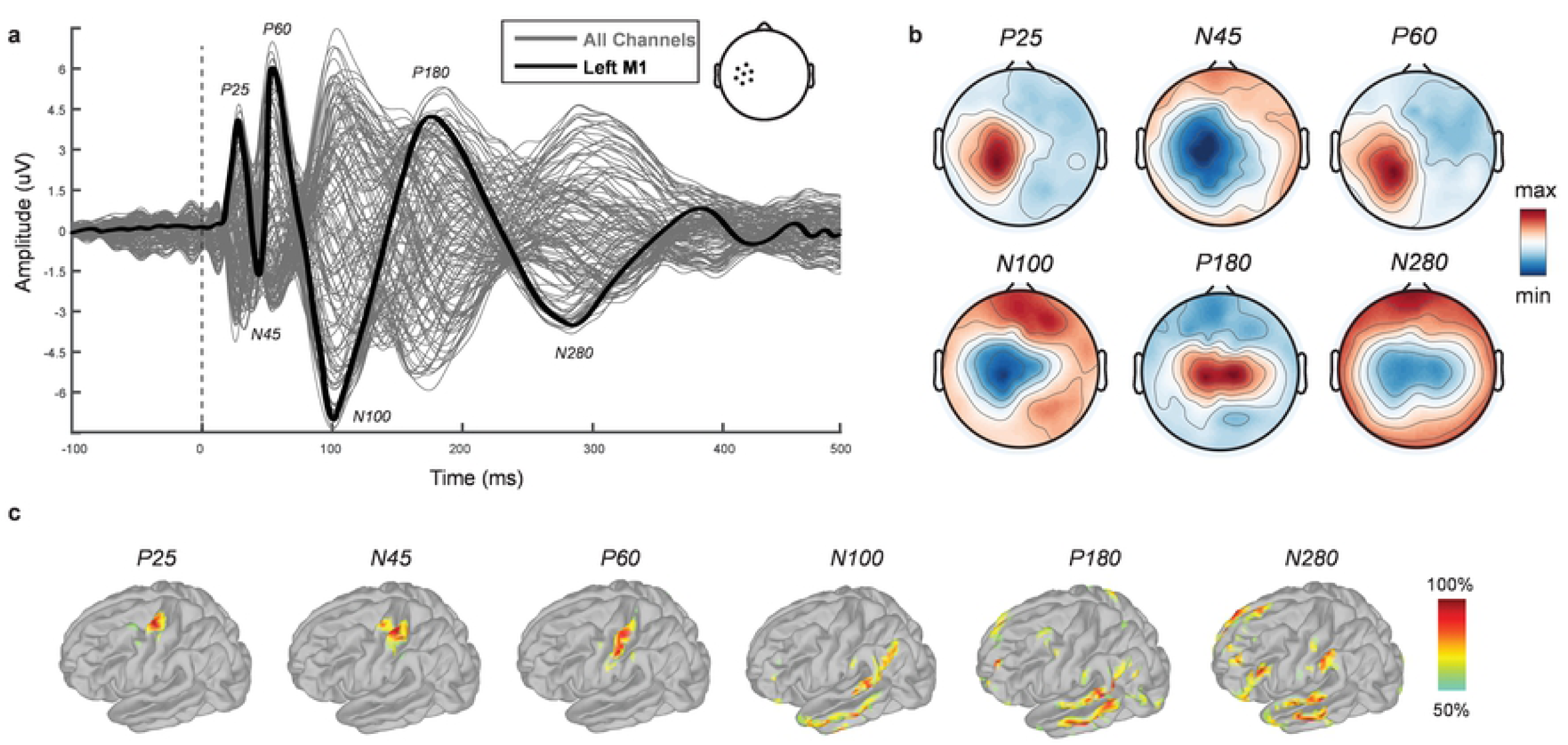
Cortical reactivity from single-pulse TMS. Time and spatial representations of TEPs in sensor and source space. (a) Butterfly plot of TEPs for all EEG channels (gray lines, 128 channels) and averaged TEP over the sensorimotor area (thick black line, 7 EEG channels). The averaged EEG channels are marked in the drawing of the scalp next to the legend. Each TEP component is referred to as P25, N45, P60, N100, P180, and N280, respectively. (b) Topographical distribution of each TEP component on the scalp. Red and blue indicate maximum and minimum EEG amplitude at each time point, respectively. (c) Source localization of each TEP component on the cortex. At each time point, cortical activation was auto-scaled and thresholded at 50% to highlight maximum cortical activation elicited by single-pulse TMS.

To investigate these TEP components’ spatial distribution on the scalp, we computed topographical distributions at each TEP time point (Fig. 1b). We found that the left sensorimotor area was activated predominantly up to 60ms (P25, N45, and P60). After P60, the centroid of activation drifted towards the midline until it centered entirely at 280ms (N100, P180, and N280).

As the sensor-space representation captures the summed cortical activity on the scalp, we next localized the TEPs on the cortex (source space). We first localized the TEPs to individual cortex models (15000 voxels) for each participant and then projected the localized TEPs to a template cortex model (15000 voxels, FsAverage) for group analysis and computed the grand-averaged TEP (5255 epochs). For each component depicted in sensor space (Fig. 2b), we projected the grand-averaged TEP onto the template cortex model (Fig. 2c). We found that P25, the earliest TEP component, was localized to the hand area of the left M1 (TMS target, Fig. 1b). N45 showed activation that spread between the M1 and the primary somatosensory cortex (Fig. 2c, second column). Next, P60 was localized to primary somatosensory cortex (Fig. 2c, third column). In contrast, the N100 and P180 peaks largely were localized to the auditory cortex and reflected the N100-P180 auditory complex[6,7]. The final TEP component (N280) also was localized to the auditory cortex, but exhibited additional activation in the frontal cortex. These findings demonstrate that single-pulse TMS on the hand area of the M1 elicits multiple TEP components and that the earliest (P25) reflects genuine cortical reactivity to TMS. We hypothesized from this finding that the N45 and P60 reflect the afferent signal from the corticospinal tract attributable to hand-muscle twitches. In contrast, the later TEP components (N100, P180, and N280) may reflect auditory processing of the coil’s clicking sound.

### Cortical reactivity and corticospinal response

Having identified cortical reactivity by single-pulse TMS in the hand area of the left M1 and evidence of an afferent signal from the primary somatosensory cortex, we next investigated how each TEP component was associated with the TMS-induced corticospinal response measured by MEPs. We averaged the TEPs and MEPs before tDCS application for each session and obtained 54 averaged TEPs and MEPs (3 sessions, 18 participants). We extracted the six TEP components (peaks) for each participant and performed correlation analyses using the Pearson correlation between the MEPs and each TEP component at each EEG channel. We found positive correlation clusters for the P25 (9 EEG channels) and P60 (6 EEG channels), and a negative cluster for the N45 (5 EEG hannels) in the left sensorimotor area (Fig. 3a, top row, *r*-value topographical maps). The black dots in the topographical maps indicate significant EEG channels (*p*<0.05). In contrast, we found no significant cluster for the N100, P180, and N280 (Fig. 3a, bottom row, *p*>0.05).

**Fig. 3.**
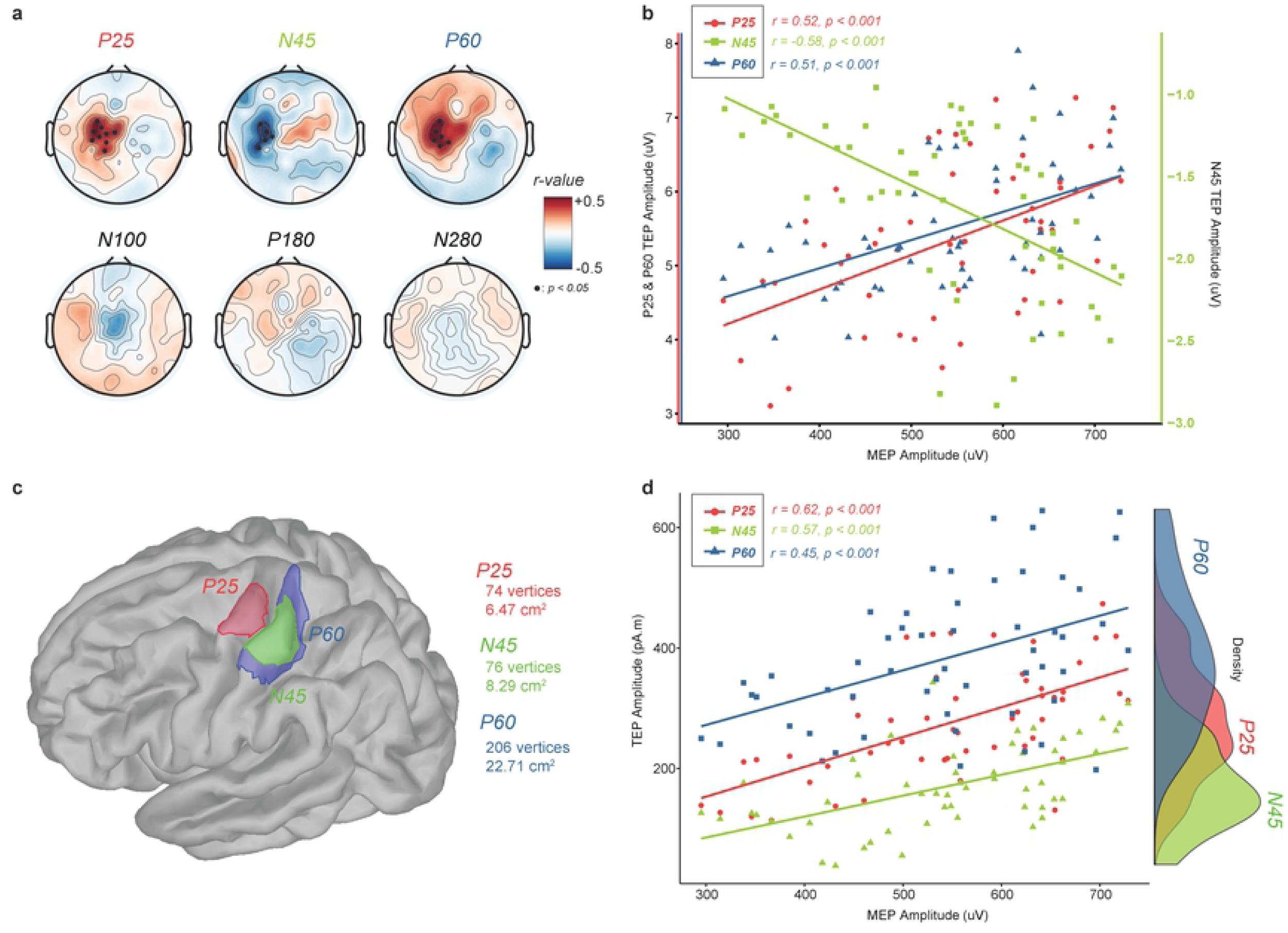
Cortical reactivity and corticospinal response. Correlation between cortical reactivity and corticospinal response. (a) Topographical distributions of correlations between each TEP component (P25, N45, P60, N100, P180, and N280) and MEPs. Black dots in topographical maps indicate significant EEG channels (*p*<0.05). P25, N45, and P60 were correlated significantly with MEPs in the left sensorimotor area, while no significant relation was found for the N100, P180, and N280. (b) Scatter plot of the averaged significant EEG channels for the P25, N45, and P60. Note that the right, green y-axis corresponds to the N45 amplitude (negative amplitude). Significant correlations were found for the P25 (*r*=0.52, *p*<0.001), N45 (*r*=-0.58, *p*<0.001), and P60 (*r*=0.51, *p*<0.001). (c) Selection of a ROI on the template cortex model (P25: 74 voxels, 6.47cm^2^, N45: 76 voxels, 8.29cm^2^, P60: 206 voxels, 22.71cm^2^). (d) Scatter plot of the ROI for each localized TEP component with MEPs. Significant correlations are obtained for the P25 (*r*=-0.62, *p*<0.001), N45 (*r*=0.57, *p*<0.001), and P60. (*r*=-0.45, *p*<0.001). The density plot shows the way the TEP components were correlated with MEPs.

To understand the relation between cortical reactivity and the corticospinal response better, we selected the significant EEG channels for each TEP component and averaged them to obtain scatter plots with MEP amplitude (Fig. 3b, n=54 for each TEP component). As expected, we found significant positive correlations for the P25 (*r*=0.52, *p*<0.001) and P60 (*r*=0.51, *p*<0.001), and a significant negative correlation (*r*=-0.58, *p*<0.001) for the N45. Note that right green y-axis corresponds to the N45 amplitude (negative amplitude)

Next, we investigated how the localized TEP components in source space were correlated with MEPs. First, we defined a region of interest (ROI, Fig. 3c) for the TEP components (P25, N45, and P60) based on the source-localized TEPs (Fig. 2c). Using these ROIs, we performed correlation analyses between the localized TEP components and MEPs (Fig. 3d). We found significant positive correlations for the P25 (*r*=0.62, *p*<0.001), N45 (*r*=0.57, *p*<0.001), and P60 (*r*=0.45, *p*<0.001). These findings indicate that the first three TEP components (P25, N45, and P60) in both sensor and source space are correlated with MEPs at baseline. Because of their location in the sensory cortex, the N45 and P60 may reflect afferent input by hand-muscle twitches.

### Modulation of motor cortex excitability by tDCS

Having identified cortical reactivity by single-pulse TMS in both sensor and source space and verified that the evoked activity predicted the corticospinal response, we next tested causally whether cortical reactivity drove the corticospinal response using tDCS to the left M1. Previous studies have shown that tDCS modulates corticospinal excitability depending upon polarity[8,9]. We hypothesized that if the TEP components reflect genuine cortical reactivity elicited by single-pulse TMS to the M1, then tDCS to the M1 should modulate the TEP components as well as MEPs in a polarity-dependent manner. We applied three different tDCS conditions (anode, cathode, and sham) at 2mA for 10 minutes and recorded MEPs and TEPs before and after tDCS. To investigate the modulation of corticospinal excitability by tDCS, we averaged the MEPs and calculated the ratio (post/pre) for each tDCS condition. Using a linear mixed-effects model, we found a significant effect of “condition” (Fig. 4a, anode vs. cathode vs. sham, *F*_2,28_=255, *p*<0.0001), but not of “session” (the three experimental sessions’ temporal order, *F*_2,28_=0.86, *p*=0.43) or their interaction (*F*_4,28_=1.56, *p*=0.21). As hypothesized, this finding demonstrated that tDCS modulated corticospinal excitability as measured by MEPs. Thereafter, we investigated whether tDCS modulated cortical excitability. We calculated the TEPs’ local mean field power in the left sensorimotor area (averaged 7 EEG channels described previously) for the entire epoch and calculated the ratio (post/pre) for each tDCS condition. We found that the period of the TEP from 25 to 60ms differed significantly for “condition” (Fig. 4b, shaded period; linear mixed-effect model, *F*_2,28_=129, *p*<0.0001), but not for “session” (*F*_2,28_=1.12, *p*=0.34), or their interaction (*F*_4,28_=1.31, *p*=0.29). In contrast, we found no significant difference for the other TEP components across tDCS conditions (100 to 280ms, *p*>0.05).

**Fig. 4.**
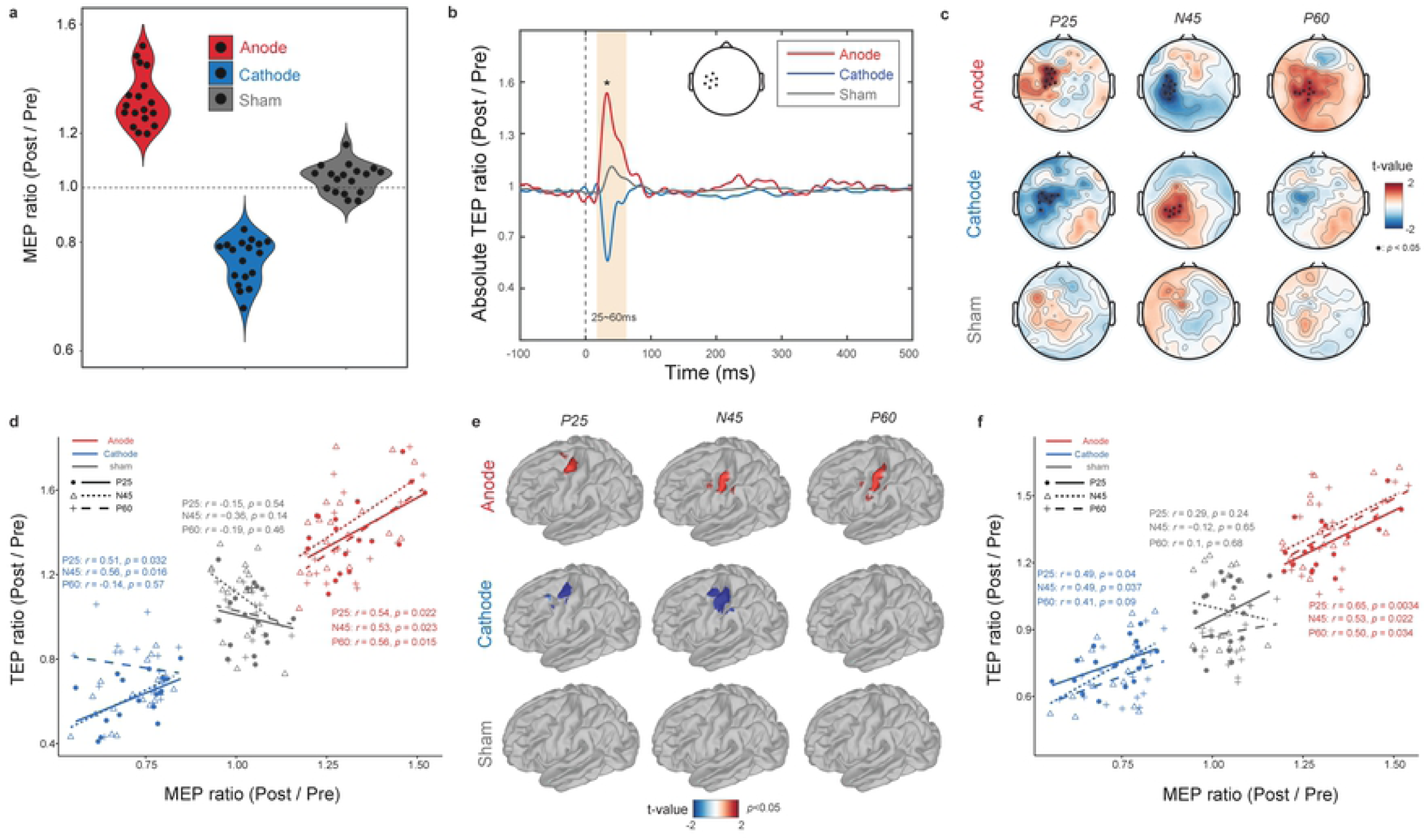
Modulation of motor cortex excitability by tDCS. tDCS modulates corticospinal and cortical excitability. (a) The ratio (post/pre) MEPs by tDCS conditions (red: anode over the left M1, blue: cathode over the left M1, gray: sham). (b) The ratio (post/pre) of absolute TEPs as a function of time for each tDCS condition. Shaded time window (25 to 60ms) differed significantly across tDCS conditions (*p*<0.0001). (c) Topographical distributions of each TEP component (*t*-value maps). A non-parametric cluster-based permutation test was performed. Black dots in the topographies indicate significant EEG channels (*p*<0.05). (d) Scatter plot of modulated corticospinal and cortical excitability in sensor space for each tDCS condition (color-coded). The markers’ shape indicates each TEP component (P25: circle, N45: triangle, P60: crosshair). Regression line to each TEP component’s scatter plot (P25: continuous line, N45: short-interval dash line, P60: long-interval dashed line). (e) Modulation of localized TEP components in source space (non-parametric cluster-based permutation test, n=1000). *t*-value maps are presented (*p*<0.05). Each row indicates tDCS conditions (anode, cathode, and sham). Each column corresponds to a TEP component (P25, N45, and P60, respectively). (f) Scatter plot between modulated corticospinal and cortical excitability in source space. Color-coded lines indicate tDCS conditions (red: anode, blue: cathode, gray: sham). The dots’ shape indicates each TEP component (P25: circle, N45: triangle, P60: crosshair). Regression line to each TEP component’s scatter plot (P25: continuous line, N45: short-interval dash line, P60: long-interval dashed line).

To investigate the modulated TEPs’ spatial representation for each tDCS condition, we next computed topographical distributions for the P25, N45, and P60. We found that the left sensorimotor area for the P25, N45, and P60 differed significantly in the anodal tDCS condition (Fig. 4c, top row, *t*-value topographical distributions, non-parametric cluster-based permutation test, n=1,000; see Supplementary Fig. 2a for the N100, P180, and N280). Black dots in each topography indicate significant EEG channels (*p*<0.05). Anodal tDCS amplified the magnitude of TEP components in the consistent direction. In the cathodal tDCS condition, we found that the sensorimotor area differed significantly for the P25 and N45, but not for the P60 (Fig. 4c, middle row; see Supplementary Fig. 2a for the N100, P180, and N280). Cathodal tDCS attenuated the magnitude of TEP components that contained M1 activation. In the sham tDCS condition, we found no significant EEG channels for the P25, N45, or P60 (Fig. 4c, bottom row; see Supplementary Fig. 2a for the N100, P180, and N280).

We then performed correlation analyses to investigate whether tDCS modulated TEPs (cortical excitability) and MEPs (corticospinal excitability) similarly across participants (Fig. 4d, scatter plot). We found significant positive correlations in the anodal tDCS condition for the P25 (*r*=0.54, *p*=0.022), N45 (*r*=0.53, *p*=0.023), and P60 (*r*=0.56, *p*=0.015), and in the cathodal tDCS condition, we found significant positive correlations for the P25 (*r*=0.51, *p*=0.032) and N45 (*r*=0.56, *p*=0.016), but not for the P60 (*r*=-0.14, *p*=0.57). We found no significant correlation in the sham tDCS condition for the P25 (*r*=-0.15, *p*=0.54), N45 (*r*=-0.36, *p*=0.14), or P60 (*r*=-0.19, *p*=0.46). Thus, the amplification or attenuation of cortical excitability measured in sensor space was consistent with the modulation of corticospinal excitability. Anodal tDCS amplified early TEP components and the degree of amplification predicted an increase in MEP amplitude, while cathodal tDCS attenuated early TEP components, which predicted a decrease in MEP amplitude.

Then, we investigated how tDCS modulated the localized TEP components by contrasting source-localized TEPs before and after tDCS. For group-level statistical tests, we projected the TEPs from individual cortex models to the template cortex model (15000 voxels). We found that the P25 differed significantly on the hand area of the left M1 (Fig. 4e, first column, non-parametric cluster-based permutation test, n=1000, *p*<0.05) in the anodal and cathodal tDCS conditions. The N45 and P60 also were modulated after anodal tDCS, but the modulation was localized in the primary somatosensory cortex (Fig. 4e, first row, second and third columns). After cathodal tDCS, the N45 was modulated significantly in the primary somatosensory cortex (Fig. 4e, second row, second column), but the P60 did not differ significantly (Fig. 4e, second row, third column). We found no such significant differences in the sham tDCS condition (Fig. 4e, third row). Similarly, we found no statistical difference for the N100, P180, and N280 in all tDCS conditions (Supplementary Fig. 2b). These findings indicate that tDCS modulates localized cortical reactivity by single-pulse TMS in the early TEP components.

We then performed correlation analyses to investigate how the modulation of localized TEPs (cortical excitability) were correlated with the modulated MEPs (corticospinal excitability) across participants (Fig. 4f). We chose the ROI on the cortex model (Fig. 3c) for each localized TEP component, and found significant positive correlations in the anodal tDCS condition for the P25 (*r*=0.65, *p*=0.0034), N45 (*r*=53, *p*=0.022), and P60 (*r*=0.50, *p*=0.034). In the cathodal tDCS condition, we found significant positive correlations for the P25 (*r*=0.49, *p*=0.04) and N45 (*r*=0.49, *p*=0.037), but not for the P60 (*r*=0.41, *p*=0.09). We found no significant correlations in the sham tDCS condition for the P25 (*r*=0.29, *p*=0.24), N45 (*r*=-0.12, *p*=0.65), and P60 (*r*=0.1, *p*=0.68). These findings support a model in which tDCS selectively modulates the localized TEP components (cortical excitability) that drive the corticospinal response.

### Modulation of pre-stimulus mu-rhythm by tDCS

Our results showed how tDCS modulated corticospinal and cortical excitability in a targeted and robust manner. These differences in response to TMS suggested that tDCS altered the state of the targeted network overall. Thus, we investigated next how tDCS modulated the network’s excitability and its activity’s oscillatory structure. We computed time-frequency representations for the entire epoch (−200 to 500ms) and performed non-parametric cluster-based permutations between before and after tDCS. We found that anodal tDCS increased the pre-stimulus mu-rhythm significantly (Fig. 5a, first row, *t*-value time-frequency map, clustered region); the increased mu-rhythm was located in the left sensorimotor area (inset, topographical distribution, black dots indicate significant EEG channels, *p*<0.05). In contrast, we found that cathodal tDCS decreased the pre-stimulus mu-rhythm significantly (Fig. 5a, second row, *t*-value time-frequency map, clustered region) as well as post-stimulus mu-rhythm around 250ms; the decreased mu-rhythm was located in the left sensorimotor area (topographical distribution, black dots indicate significant EEG channels, *p*<0.05). In the sham tDCS condition, we found no significant difference in the time-frequency map (Fig. 5a, third row, *t*-value time-frequency map) and topographical distribution (no significant EEG channel).

**Fig. 5.**
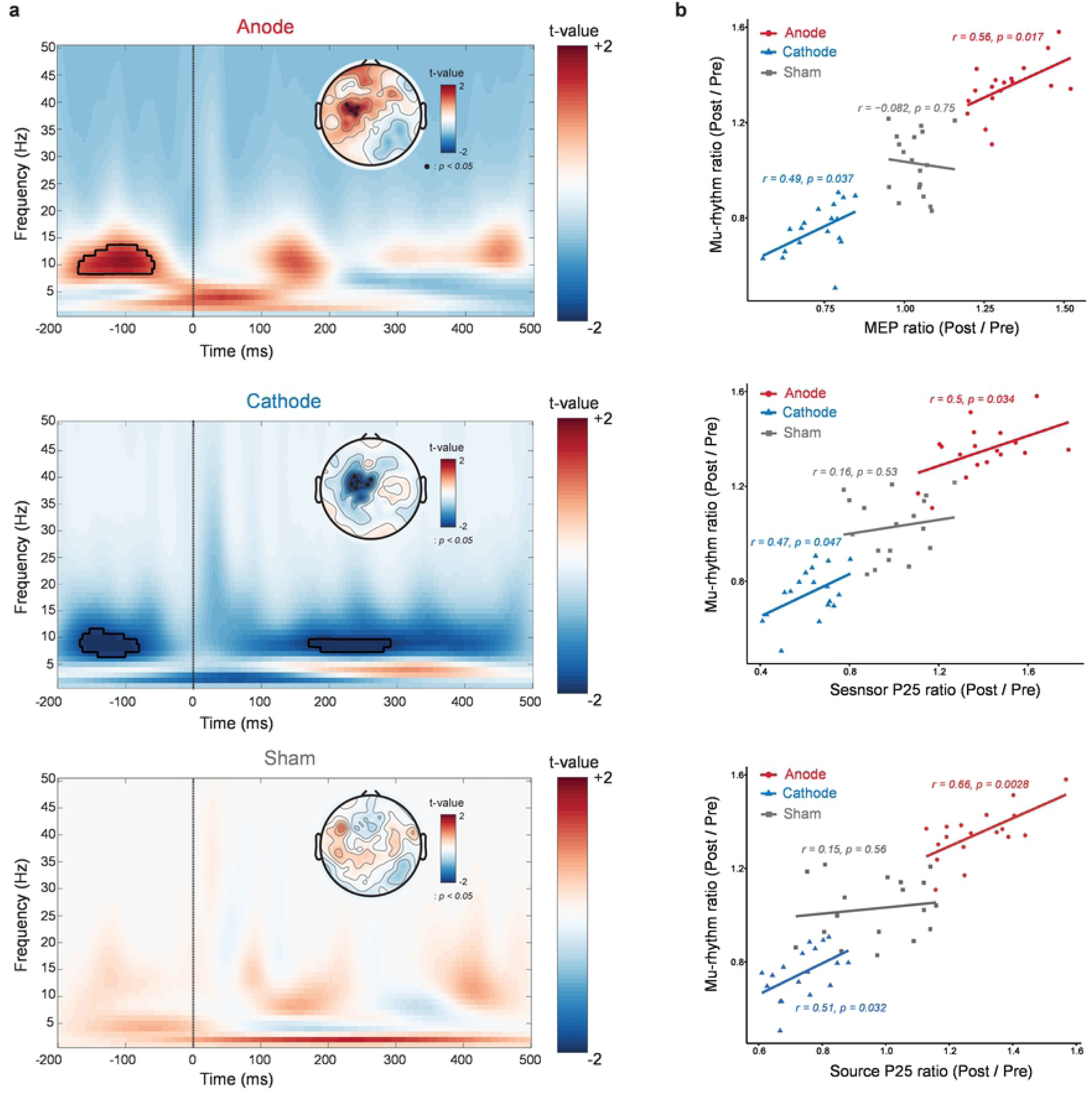
Modulation of pre-stimulus mu-rhythm by tDCS. tDCS modulates pre-stimulus mu-rhythm. (a) time-frequency maps of modulated oscillations and topographical distributions for anodal (top), cathodal (middle), and sham (bottom) tDCS conditions. Clustered region in time-frequency maps indicates significant modulation by tDCS (non-parametric permutation test, n=1000) and black dots in topographical distributions indicate significant EEG channels (*p*<0.05). (b) Scatter plots of the ratio of the pre-stimulus mu-rhythm to the ratio of MEP (top), P25 in sensor space (middle), and P25 in source space (bottom). Each dot indicates a participant and the color code indicates the tDCS conditions (red: anodal, blue: cathodal, gray: sham). Correlation coefficients (*r*-values) and *p*-values are presented.

Thereafter, we investigated the relation between the pre-stimulus oscillatory modulation and the modulation of corticospinal and cortical excitability. Correlations were calculated between the modulated pre-stimulus mu-rhythm and both MEPs and P25 TEP component in sensor and source space for each participant (Fig. 5b). We found that the ratio of the pre-stimulus mu-rhythm (post/pre to tDCS) was correlated with the ratio of MEP (post/pre to tDCS) for anodal (*r*=0.56, *p*=0.017) and cathodal tDCS (*r*=0.49, *p*=0.037), but not for sham tDCS (Fig. 5b, first row, *r*=-0.08, *p*=0.75). We also found that the ratio of the pre-stimulus mu-rhythm was correlated with the ratio of the P25 in sensor space for anodal tDCS (*r*=0.50, *p*=0.034) and cathodal tDCS (r=0.47, p=0.047), but not for sham tDCS (*r*=0.16, *p*=0.53). Similarly, we found that the ratio of the P25 in source space (ROI-based) was correlated with the ratio of the mu-rhythm for anodal tDCS (*r*=0.66, *p*=0.0028) and cathodal tDCS (*r*=0.51, *p*=0.032), but not for sham tDCS (*r*=0.15, *p*=0.56). These results show that tDCS modulates the pre-stimulus mu-rhythm and that this modulation of network oscillations altered corticospinal and cortical excitability.

## Discussion

TMS-EEG studies have gained attention recently, as they may provide important insights into disease processes in the central nervous system, as well as a mechanistic understanding of the way clinical TMS paradigms engage brain networks[2,12]. However, there is a central controversy about whether the TEP reflects genuine cortical reactivity to TMS or whether it consists of reactivity from peripherally- and auditory-evoked potentials[3– 5]. Our study addressed this controversy directly through a unique combination of brain stimulation and imaging methods. We used sophisticated procedures to attenuate the peripheral and auditory confounds generated and performed source localization with high-density EEG data, structural MR images, and digitized EEG electrode locations to obtain a high spatial resolution picture of cortical reactivity. We observed six TEP components and found that the P25, the earliest TEP component, was localized to the stimulated cortical area (the left M1). The following two TEP components (N45 and P60) largely were localized to the primary somatosensory cortex, which represent afferent input by hand-muscle twitches. The remaining TEP components (N100, P180, and N280) were localized primarily to the auditory cortex. Importantly, tDCS modulated the first two TEP components (P25 and N45) selectively depending upon polarity in our double-blind, placebo-controlled study. In addition, we found evidence that cortical reactivity played a causal role in predicting corticospinal excitability. Thus, our findings demonstrate that the early TEP reflects genuine cortical reactivity and later TEP components are associated with somatosensory and auditory processing in the brain.

A recent study that investigated neural effects at the single-cell level has shown that suprathreshold single-pulse TMS elicits a stereotyped burst of action potentials within the first 30ms (10-30ms) after TMS onset in the macaque parietal cortex[13]. Another study with human participants found that single-pulse TMS to the M1 resulted in significant differences before 60ms compared to sham TMS[14]. Consistent with these recent findings, we found that the P25 was localized to the left M1 (TMS target location), demonstrating that the P25 represents genuine cortical reactivity to single-pulse TMS to the M1. Although we were unable to obtain earlier TEP components, such as the P10[11] or P15[15] because of the TMS artifacts in our recordings, the response latency (within 30ms) is consistent with previous findings. We also observed N45 and P60 components that were localized primarily in the primary somatosensory cortex and reflected afferent input by hand-muscle twitches produced by suprathreshold TMS. We demonstrated further that these somatosensory-evoked potentials were correlated with MEP amplitude (Fig. 3b) and comparable to the conventional somatosensory evoked potentials with respect to response latency[16]. For the later TEP components, although we applied auditory masking using white noise that removed the auditory perception of TMS pulses, we obtained the typical N100-P180 auditory complex[6] by single-pulse TMS (Fig. 2a), which was localized in the auditory cortex (Fig. 2c). This phenomenon may derive from inevitable bone- and air-conducted sound from the TMS coil[7]. The amplitude of these potentials (>5uV) was comparable with the N100 amplitude in our study. Thus, we conclude overall that each TEP component single-pulse TMS elicits has a distinct network representation in the brain and the P25 represents genuine cortical reactivity from TMS to the M1.

Since the first attempt to modulate motor cortex excitability by weak direct current on the scalp[17], it has been shown consistently that tDCS modulates motor cortex excitability depending upon polarity[8,9,18–23]. We hypothesized that if a TEP elicited by single-pulse TMS on the M1 is genuine motor-related cortical reactivity, then tDCS to the M1 should modulate it. We found that tDCS successfully modulated the P25 in the stimulated cortical area in a polarity-dependent manner (Fig. 4e). tDCS also modulated the N45 in the same manner, but only anodal tDCS modulated the P60. Consistent with the findings for the P60, the relation between changes in MEP and P60 amplitude was not significant in both the sensor (*r*=-0.14, *p*=0.57) and source (*r*=0.41, *p*=0.09) space. We assume that this unexpected finding might be caused by the reduction of post-stimulus mu-rhythm (around 200 to 300ms after onset) by cathodal tDCS (Fig. 5b, second row, time-frequency *t*-value map). We hypothesized that tDCS could modulate only the pre-stimulus mu-rhythm, but cathodal tDCS actually reduced the post-stimulus mu-rhythm, which was not found in the anodal tDCS condition. This inconsistency in modulation of cortical reactivity should be investigated in the future. While we adopted the conventional M1-SO montage for tDCS, which uses two stimulation electrodes (5×7 rectangular electrodes, one on the motor area and another on the supraorbital area) to modulate motor cortex excitability, a recent study used a 4×1 montage that consisted of smaller, ring-shaped electrodes (referred to as high-definition tDCS, HD-tDCS) that was introduced to increase the focality of induced electric field[24]. One study[19] compared the effect of modulating motor cortex excitability between the two montages and found that both have a comparable effect in modulating excitability. In our study, we used the M1-SO montage with two smaller electrodes (5×5cm, 25cm^2^) to increase efficacy via a greater current intensity[20]. We performed electric field modeling with structural MR images and confirmed that the induced electric field is comparable to that in previous tDCS studies (Fig. 1c). As an exploratory analysis, we investigated how the induced electric field in the target stimulation area is related to MEP changes (Supplementary Figure 3) inspired by a study[25] that found that the intensity of the electric field in the primary motor cortex can explain inter-individual variability in MEP. However, we found no relation between them; thus, this finding may suggest that more factors, such as phase-dependent excitability, could have affected the motor cortex excitability modulation in our data[26].

The corticospinal response (measured by MEP) elicited by single-pulse TMS on M1 varies between trials[27–30]. Recent studies have shown that this variability is associated with neural oscillation power[31–37], phase[26,38,39], or their interaction[40], although one study failed to replicate these findings[41]. In our study, we showed that pre-stimulus mu-rhythm oscillatory power was correlated with the modulation of cortical and corticospinal excitability (Fig. 5b). This finding indicates that tDCS modulates oscillatory power and thereby, the modulated oscillatory power causes the modulation of cortical and corticospinal excitability. Consistent with this causal role of oscillatory power, a previous study showed that anodal tDCS increased neural oscillatory power and altered functional connectivity in a non-human primate model[42]. Importantly, recent TMS-EEG studies have found that pre-stimulus oscillatory power was correlated positively with MEP amplitude[35,37]. Together, our findings may represent the causal role of oscillatory power in motor cortex excitability.

As with any scientific investigation, this study has limitations. First, we were unable to study the earlier TEP components at 10[11] or 15ms[15] because of TMS pulse artifacts. We used a TMS-compatible EEG amplifier (NetAmps 410, Philips Neuro Inc.), but we observed that the TMS pulse artifact lasted up to 20ms in raw EEG traces (Supplementary Figure 5). Although we demonstrated that the P25 was localized on the hand area of the M1, future investigations of the earlier TEP components should be considered with an EEG amplifier that has a faster recovery period. Second, although we demonstrated that the P25 reflects genuine cortical reactivity from TMS to M1, we did not show TEP dynamics of single-pulse TMS to other brain regions, such as the dorsolateral prefrontal cortex, which is the main target in the treatment of depression[43,44]. Previous studies have shown that TEPs exhibit different dynamics[45–48] thus the comparison of stimulation to different cortex regions should be investigated in the future to confirm our findings.

The event-related potential (ERP), which is an evoked EEG potential in response to an external stimulus, has been studied well over the past several decades[49]. Each ERP component represents specific processing in the brain. For example, the P300, a positive peak potential at approximately 300 milliseconds, reflects cognitive processing[50], while the N170, a negative peak potential at approximately 170 milliseconds, is a face-recognition ERP component over the ventral area of the visual cortex[51]. However, in the field of TMS-EEG, few efforts have been made to determine how each TEP component is associated with specific sensory processing, and the underlying mechanism remains unclear. As the number of studies, used TMS as a treatment tool, has increased tremendously in recent years, understanding of how the brain responds to TMS is imperative to both the research and clinical fields. Without the ability to interpret TEP components appropriately, the rational design and subsequent optimization of network-based treatment strategies with non-invasive brain stimulation is jeopardized. In our study, thus, we sought to bridge the intellectual gap and it may have a large impact on the field.

In summary, we demonstrated that the early TEP reflects genuine cortical reactivity elicited by single-pulse TMS. We identified each TEP component in sensor and source space and used tDCS to modulate the TEP components successfully in a polarity-dependent manner, and found that the modulation of the pre-stimulus mu-rhythm by tDCS caused the modulation of excitability. Further, we found that the TEP components (cortical excitability) were correlated significantly with MEP amplitude (corticospinal excitability). These findings suggest that each TEP component plays a distinct role in specific sensory processing in the brain.

## Methods

### Study design

We performed a crossover, double-blind, sham-controlled study with three tDCS conditions (anodal, cathodal, and sham tDCS) at the University of North Carolina at Chapel Hill, which the Biomedical Institutional Review Board at the university approved. The study protocol was registered before participants were recruited (ClinicalTrials.gov, NCT03481309). We recruited 19 healthy, right-handed, male participants free of any neurological disorders. All participants provided written informed consent before participation. After telephone screening to assess their eligibility for the study, structural MR images (T1-weighted) were obtained using a 3T-MRI scanner (Magnetom Prisma, Siemens AG, Berlin, Germany) at the University of North Carolina Biomedical Research Imaging Center. One of the participants dropped out of the study because of perceived scalp discomfort attributable to TMS. All remaining participants completed the three tDCS sessions, in which the order of the conditions was distributed equally (three participants per each tDCS order). There was at least a 3-day interval between the sessions to minimize any (theoretical) long-lasting effects of tDCS. Each session consisted of the following procedures (Fig. 1): determination of RMT, EEG, and MEP recordings with 100 single-pulse TMS (5 minutes, 120% relative to RMT), tDCS (11 minutes, 2mA), EEG and MEP recordings with 100 single-pulse TMS (5 minutes), and digitization of EEG electrode locations using a stereo-camera tracking digitizer (GeoScan Sensor Digitization Device, Philips Neuro Inc., Eugene, OR).

### EEG and MEP recordings with TMS

Based on the structural MR images, we performed brain segmentation and determined an initial target location (hand area on the left M1) using a frameless neuronavigation system (Localite GmbH., Sankt Augustin, Germany). According to the initial target location, a figure-of-eight coil (C-B60, MagVenture Inc., Farum, Denmark) was placed tangentially on the scalp with the handle pointing backwards and laterally at 45 degree from the mid-sagittal line. Participants were seated in a comfortable armchair (TMS chair) with their hands positioned on the armrests. Three EMG electrodes (15×21mm, Ambu Neuroline 700, Ambu Inc., Columbia, MD) were placed in a tendon-belly arrangement on the first dorsal interosseous muscle (active and reference EMG electrodes) and the styloid process of the ulna on the right hand (ground EMG electrode). Biphasic single-pulse TMS was applied on the initial location and the location was adjusted to obtain the highest MEP at the same intensity. MEP traces were visualized in a built-in display on the TMS device (MagPro X100, MagVenture Inc., Farum, Denmark). The RMT was defined by the minimum TMS intensity required to evoke MEPs of at least 50 uV in 50% of 5 to 10 consecutive trials[52]. The left motor hotspot (hand area on the M1) was determined at this step. We used the Physio 16 input box (Philips Neuro Inc., Eugene, OR) connected to the EEG amplifier to record MEPs. This configuration allowed us to record MEP and EEG data on the same amplifier. We used a TMS-compatible EEG system with a 128-channel net (Philips Neuro Inc., Eugene, OR) at a sampling rate of 1kHz. Channel Cz and one channel between Cz and Pz were used as a reference and ground, respectively. Participants wore air-conducting earphone tubes (ER-3C, Etymotic Research Inc., Elk Grove Village, IL) with white-noise masking to attenuate auditory evoked potentials[11]. We also applied a thin layer underneath the TMS coil to attenuate peripherally-evoked potentials. We applied 100 single-pulse TMS pulses (120% intensity relative to RMT) with a jittered inter-trial interval between 2 and 3 seconds to minimize any anticipatory effect. All TMS pulse locations were tracked in real-time using the neuronavigation system and saved for verification of stimulation on the left motor hotspot. The EEG and MEP recording procedures were performed both before and after tDCS.

### Transcranial direct current stimulation (tDCS)

We applied two carbon-silicone electrodes (5×5cm) to the scalp with Ten20 conductive paste (Bio-Medical Instruments, Clinton Township, MI) and used the XCSITE 100 stimulator (Pulvinar Neuro LLC, Chapel Hill, NC). The stimulator does not display any information about the stimulation conditions (verum or sham). The two electrodes were placed at the location of the left motor hotspot (determined by RMT) and the right supra-orbital area (Fp2 EEG location based on the 10-20 international coordinate system). In anodal tDCS, we delivered 11 minutes and +2mA of constant current, including 60 seconds of ramp-up and - down (10 minutes of +2mA constant current). In cathodal tDCS, we delivered 11 minutes of stimulation, including −2mA of constant current and 60 seconds of ramp-up and -down (10 minutes of −2mA constant current). In sham tDCS, we delivered 30 seconds of +2mA constant current with 60 seconds of ramp-up and -down. The choice of such an “active” sham is an established strategy to enhance blinding the participants to the stimulation conditions[53]. After the trials, all participants were asked to fill out a questionnaire indicating whether they received electrical stimulation or not (Yes or No) and side-effect questionnaires (Supplementary Figure 6). We found no significant differences in the side-effect questionnaires among the tDCS conditions.

## Data analysis

### MEP and EEG data analysis

Offline data processing was performed with custom-built scripts in MATLAB (R2015b, Mathworks Inc., Natick, MA) and the EEGLAB toolbox[54]. The MEP data collected were inspected visually and epochs that had less than 50uV MEP were removed (4.4±7.2 of 100 epochs). MEP data were averaged for each condition (before and after TMS) and the ratio (pre/prost) was calculated. The ratio at each session represents modulation of MEPs by tDCS. To analyze the EEG data by single-pulse TMS, we identified the TMS onset and TMS-induced artifacts (−10 to 20ms to the TMS onset) first. This artifact time period was replaced by a value selected randomly from a Gaussian distribution made by the standard deviation and mean of a reference period (−50 to −10ms to the TMS onset)[55]. Second, the data were band-pass filtered from 1 to 50Hz. Third, the data were preprocessed by an artifact subspace reconstruction algorithm[56] to identify high-variance data epochs and reconstruct missing data. Fourth, bad channels that were found in the previous step were interpolated and common average referencing was performed. Thereafter, infomax independent component analysis (ICA)[57] was performed to remove eye blinking, eye movement, muscle activity, and heartbeat artifacts. All ICA components were inspected visually and noise components were selected manually for rejection. The selection of ICA components were verified by the ICLabel classification[58]. The preprocessed EEG data were epoched from −100 to 500ms to the respective TMS onset. Each epoch was inspected visually and noisy epochs were removed (3.7±6.1 of 100 epochs). We found no significant difference between the three conditions in the epochs rejected (one-way ANOVA, *F*_2,51_=0.72, *p*=0.49). To obtain a grand-averaged TEP for each channel, we averaged 5255 epochs after epoch rejection across participants and conditions (before tDCS) as a function of time (−100 to 500ms). We used the Morlet wavelet transform (7 cycles) with a frequency resolution of 1Hz and temporal resolution of 1ms to compute time-frequency maps of the entire epoch (−200 to 500ms) for each channel. The power in the time-frequency maps was obtained and was used for statistical tests across tDCS conditions.

### EEG source localization

After obtaining structural MR images for each participant, we performed skull stripping, gray-white matter segmentation, reconstruction of cortical surface models (gray-white boundary surface and pial surface), and labeled regions on the cortex using FreeSurfer 5.3[59]. Preprocessed and segmented MR images were imported in the BrainStorm toolbox[60]. Three fiducial points (nasion and left/right preauricular points) and anatomical points (anterior/posterior commissure and inter-hemispheric point) were defined on the MR images. We built a scalp model consisting of 10000 vertices from the MR images and co-registered it with digitized EEG electrodes locations for each session. During this step, we confirmed that all scalp EEG electrodes were projected properly on the scalp model. We used the boundary element method (BEM) with OpenMEEG[61,62] to compute the lead field matrix (forward modeling). The forward model consisted of 9808 vertices for the scalp (conductivity: 1), 1922 vertices for the skull (conductivity: 0.012), and 1922 vertices for the brain (conductivity: 1). After obtaining the forward model for each session and participant, we used the linearly constrained minimum variance beamformer[63] to solve the ill-posed inverse problem (inverse modeling). We projected scalp EEG signals to the cortex model consisting of 15000 voxels. We averaged all projected source activity on the individual cortex model across trials and projected it onto the template cortex model (FSAverage, 15000 voxels) for group-level analysis[64].

### Statistical testing

We used the linear mixed-effects model in R (R Foundation for Statistical Computing, Vienna, Austria) to investigate modulation of cortical and corticospinal excitability with the fixed factors of “tDCS condition” (anode, cathode, and sham) and “session” (sessions 1, 2, and 3), with the random factor, “participant”. The dependent variables were the ratio of averaged MEPs and the ratio of averaged TEPs over EEG channels.

To calculate the spatio-temporal statistical significance for TEPs in both sensor and source space for each tDCS condition, we used a non-parametric cluster-based permutation test[65] to address the multiple comparison problem of high-density EEG. First, *t*-tests were conducted for each channel and time point across participants between before and after tDCS for each tDCS condition. We then constructed clusters from the spatio-temporal significant *t*-value map (*p*<0.05) obtained, summed all of the positive or negative *t*-values within the clusters separately, and clustered the significant *t*-values based on spatio-temporal adjacency. The minimum size of a cluster was set to two points. A neighboring channel was defined as spatial adjacency within 4 cm[65]. For the permutation test, we shuffled all trials and divided them into two datasets. We then conducted *t*-tests for the two datasets to obtain a *t*-value map. We repeated this procedure by Monte Carlo simulation with 1000 iterations, and extracted the largest cluster from each permutation test to compare with the original dataset. Lastly, we constructed a histogram of the 1000 values of the cluster-level statistics and calculated a probability density function (PDF) to estimate cluster-level *p*-values. The input for the PDF was the cluster-level statistics from the original dataset, while the output was a *p*-value for each cluster-level statistic. The cluster-level *p*-values were corrected and approximated by this cluster-based permutation test.

## Author Contributions

S.A. and F.F. designed the study. S.A. collected and analyzed the data. S.A. and F.F. wrote the manuscript.

## Acknowledgements

The authors thank Julianna H. Prim for her assistance of tDCS electrode application, Dr. Kai Xia for providing his expertise in statistical analysis, and Dr. Sankaraleengam Alagapan, Dr. Justin Riddle, Trevor McPherson for their feedback in study design. The authors specially thank Donghyeon Kim (Neurophet Inc.,) for providing valuable feedback on electric field modeling and Dr. Zhe Charles Zhou for his work in creating randomization codes and validating tDCS conditions for double-blinding. The authors thank Dr. Justin Riddle for his work in validating tDCS waveforms after completion of the study. The authors thank Trevor McPherson, and Dr. Justin Riddle for their feedback on the manuscript. This work was supported by the National Institute of Mental Health of the National Institutes of Health under Award Numbers R01MH111889 and R01MH101547. The content is solely the responsibility of the authors and does not represent the official views of the National Institutes of Health.

## Data availability

All data, as well as analysis codes that were used to perform analyses, can be made available from the corresponding author upon reasonable request.

